# DeepBCR: Deep learning framework for cancer-type classification and binding affinity estimation using B cell receptor repertoires

**DOI:** 10.1101/731158

**Authors:** Xihao Hu, X Shirley Liu

## Abstract

We developed a deep learning framework to model the binding specificity of B-cell receptors (BCRs). The DeepBCR framework can predict the cancer type from a repertoire of BCRs and estimate the binding affinity of a single BCR. We designed a peptide encoding network that includes an amino acid encoding layer, k-mer motif layer, and immunoglobulin isotype layer, and used transfer learning to reduce parameters and over-fitting. When we applied the framework to evaluate the three commercial anti-PD1 drugs (Opdivo, Keytruda, and Libtayo), the predicted binding affinities correlate with the real affinities measured in kD values. This validates the prediction and indicates that we can use the framework to select strong antigen-specific binders.

## Introduction

B cells are able to infiltrate into the tumor and perform diverse functions in both suppressing and promoting cancer cell growth^1^. One cause of such contradictory outcomes is due to the diversity in the B cell isotypes which interact with receptors preferentially expressed on different immune cells^2^. For example, IgA antibodies reduce the anti-tumor effects in liver cancer^3^ and melanoma^4^, although the role of IgA antibodies are more complex in other cancer types^5^. On the other hand, the binding affinity of tumor infiltrating B cells is believed to be high regardless of downstream regulations, because of the wide-spread somatic hypermutations observed in tens of millions of tumor-infiltrating B-cell receptors (BCRs)^5^.

Although billions of BCRs have been sequenced in published studies^6^, they came from donors of limited clinical annotations and most of them are naive B cells. The over 30 million BCRs derived from our previous study from TCGA^5^ have rich clinical annotation and so are more likely to gain insights on the general antibody-antigen binding patterns. To understand the power of associating antibodies with antigens, we start with a simpler problem---whether we can predict cancer type from the BCR repertoire. Then, we tried to associate each BCR to individual tumor antigens with the goal of approximating the real binding affinity.

## Results and Discussions

### Tumor infiltrating B cells associate with cancer-type specific antigens in the tumor microenvironment

We chose 11 TCGA cancer types which have at least 300 patients to define the cancer-type classification problem (Fig. 1a). The B cell receptor (BCR) repertoire was collected in our previous study^5^ with an inclusion of both B cell heavy chain and light chain CDR3 regions. BCRs from the same patients were compiled together and fed into the peptide encoding network (PEN) for learning the function to output one of the 11 cancer types. We trained the networks under a wide range of parameter choices and tested the model performance on a separate test set. To evaluate the power of distinguishing a specific cancer type from the other 10 cancer types, we plotted the receiver operating characteristic (ROC) curve and calculated the area under the curve (AUC) as a metric (Fig. 1b). The model has the best power for detecting ovarian cancer (AUC=0.98) and gastric cancer (AUC=0.96) patients and the worst performance in detecting colorectal and breast cancers (AUC=0.77 for both). We further plotted the confusion matrix to investigate the causes of wrong predictions (Fig. 1c). We found that colorectal tumors were likely to be predicted as endometrial carcinoma, implying the similarities in their infiltrating B cell repertoires.

**Figure 1:**
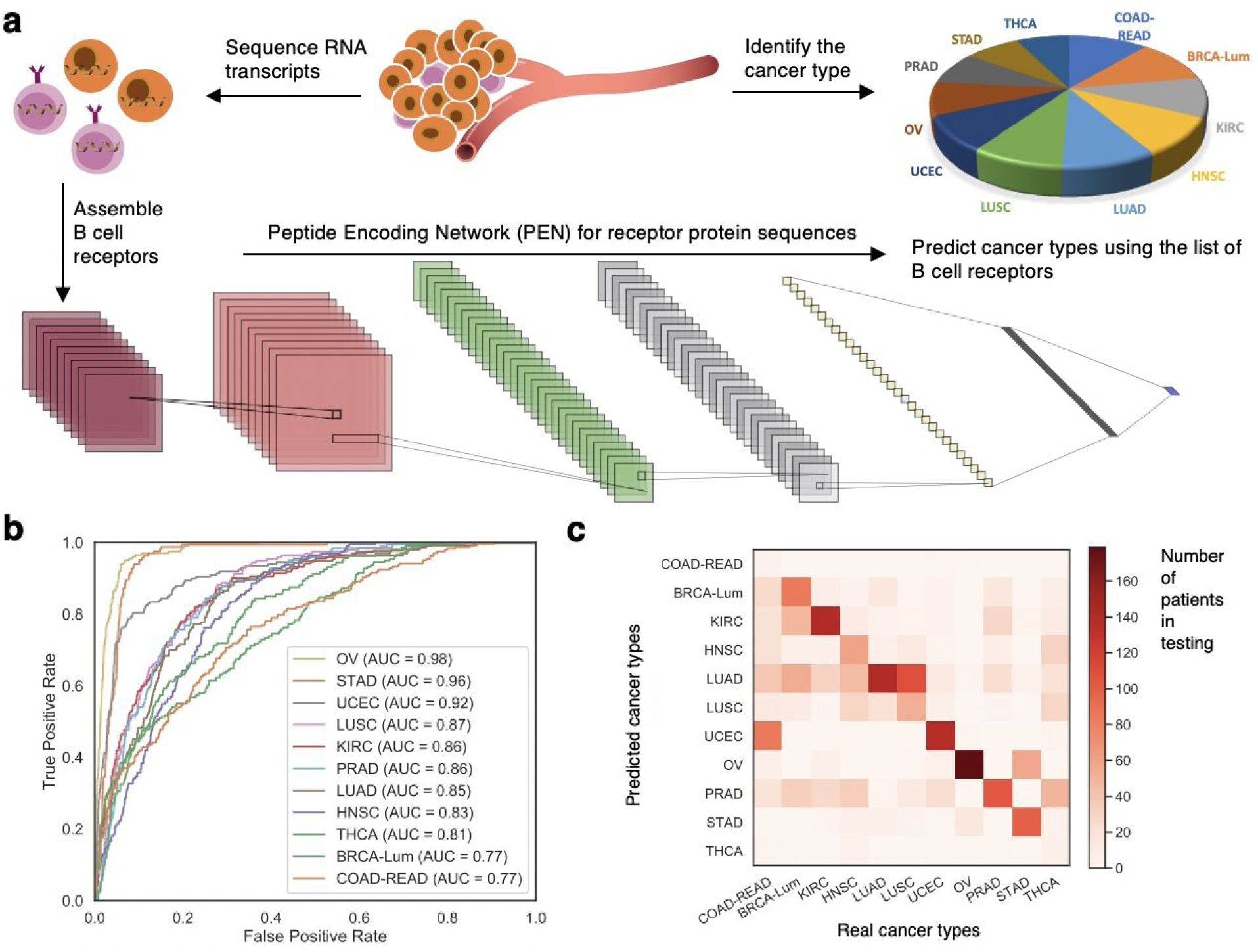
Deep neural network for the cancer type classification based on B cell receptors in the tumor microenvironment.

### Cancer-type specific B cells show similar binding specificity across tumor types

Besides modeling cancer-type specific B cells, we next modeled association of BCRs with specific tumor antigens. By adjusting the DeepBCR model using transfer learning, we found that the performance on the training and testing sets are very similar, indicating reduced risk of over-fitting (Fig. 2a). We grouped the binding targets by their gene annotations and found that tumor infiltrating B cells are more likely to bind to membrane proteins than transcription factors (Fig. 2b). We found that genes enriched in pan-cancer study also show enrichment when analyzing each cancer type separately (Fig. 2c). Pathway analyses further showed that genes associated with antigen binding are consistently high in pan-cancer and cancer-type specific studies (Fig. 2d).

**Figure 2:**
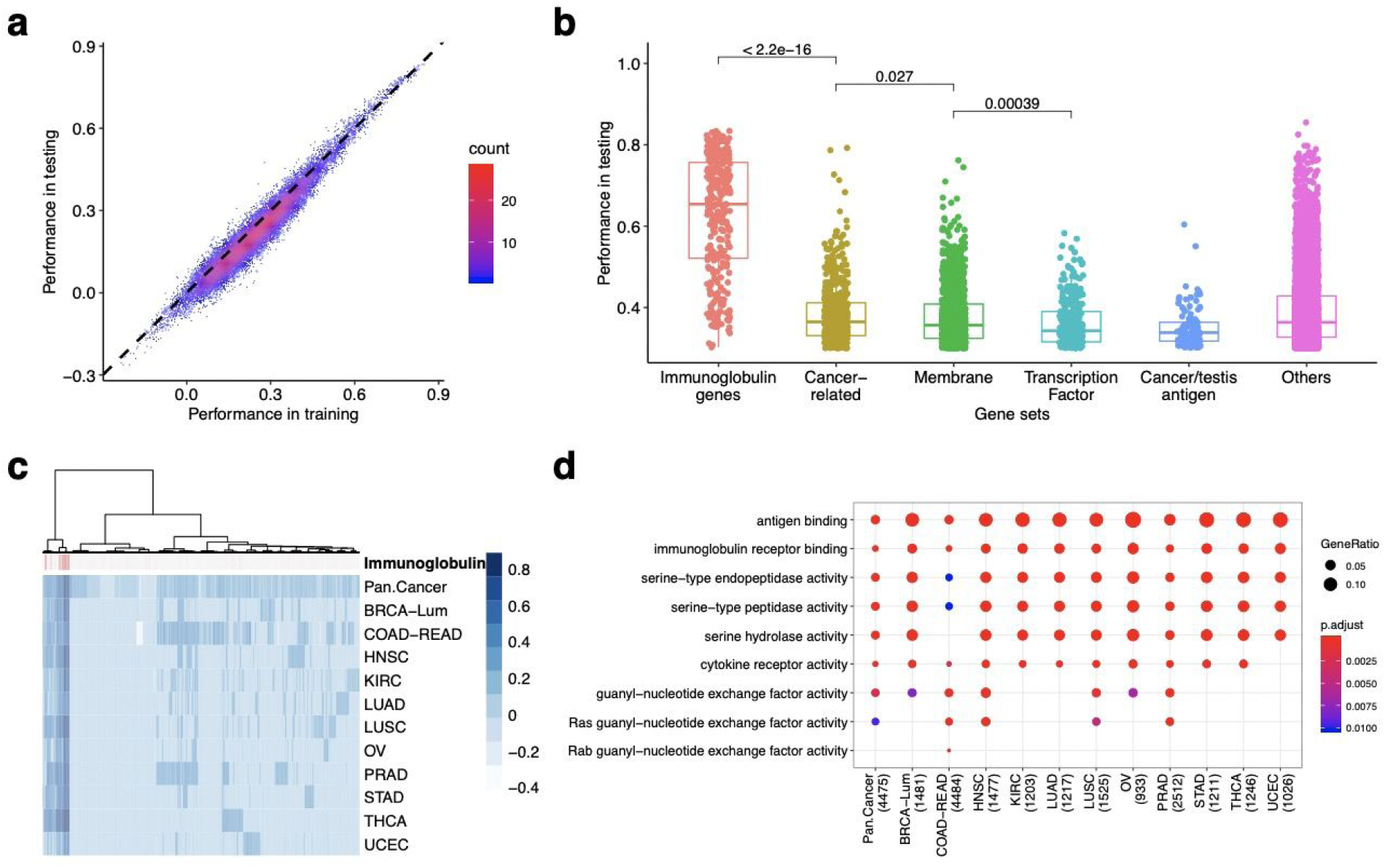
Genome-wide modeling on the antibody binding specificity.

### DeepBCR predicts the binding affinity of anti-PD1 antibody drugs based on protein sequences

We obtained the heavy and light chain CDR3 protein sequences and their binding affinity to human PD1 protein of the three commercial anti-PD1 antibody drugs from publicly disclosed information. We found that the BCR scores predicted by the DeepBCR model are negatively correlated with real binding affinity measured by kD values (Fig. 3), suggesting that the DeepBCR model predicts the binding affinity in the right direction with good association with the log kD of binding affinity. This quantitative relationship suggests that it is possible to design a new antibody with stronger binding affinity than pembrolizumab by introducing random mutations on existing antibody sequences. Our results also indicate that DeepBCR not only has the potential to predict antibody sequences against tumor antigens, eliminating the need for immunization or phage-display, but also improve binding using *in silico* affinity maturation to predict previously unobserved binders.

**Figure 3:**
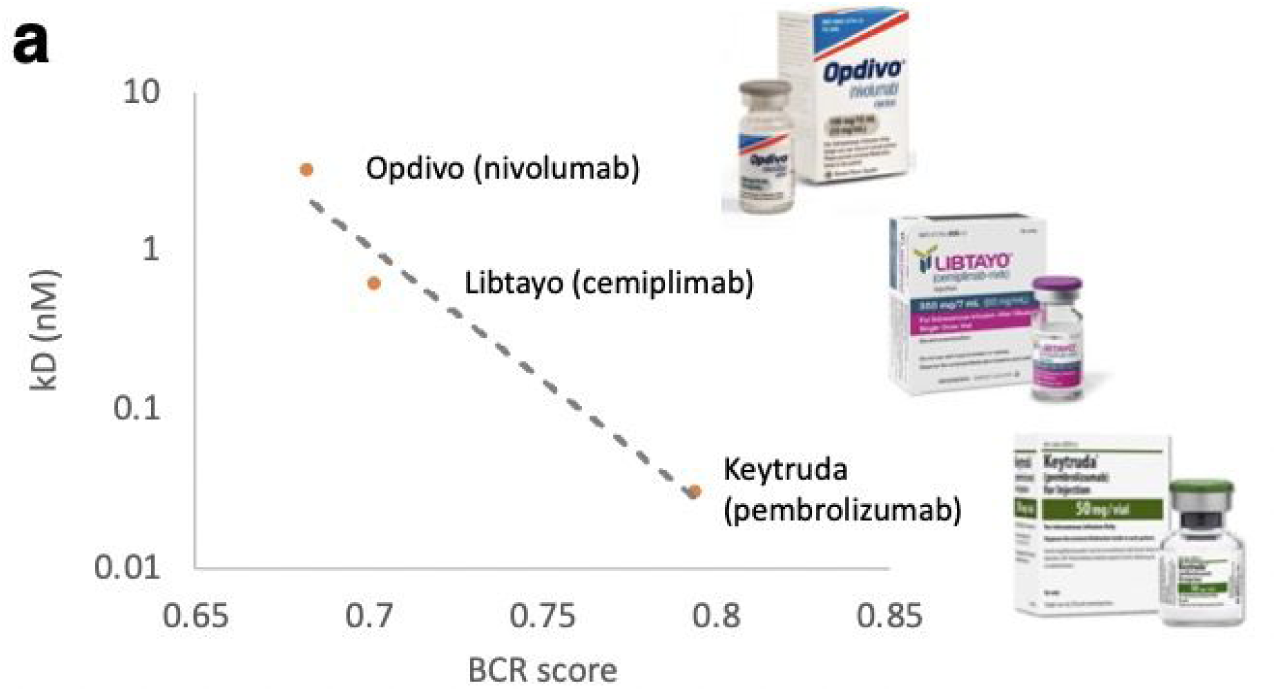
DeepBCR accurately predicts the binding affinity of commercial anti-PD1 antibody drugs.

## Conclusion

We previously developed a computational method TRUST to assemble over 30 million B cell receptors (BCRs) from TCGA bulk tumor RNA-seq data^5^. We now describe a deep learning framework, DeepBCR, to model complex antigen-antibody interactions starting from BCR sequences. We tested the model performance in associating cancer-type specific antibodies and antigen-specific antibodies, and showed the power of predicting the binding affinity of three commercial anti-PD1 antibodies drugs. Existing experimental approaches can only profile potential antibodies for a known antigen or profile potential targets for an existing antibody. In comparison, DeepBCR can model the binding specificity of many antigen-antibody pairs *in silico* with the potential of finding novel anti-cancer targets and high quality fully-human antibodies.

## Code availability

https://bitbucket.org/liulab/deepbcr

## Acknowledgement

This work was supported by the National Institute of Health U01 CA226196 and Breast Cancer Research Foundation (to X.S.L).

